# Friend or foe? Effects of host immune activation on the transient gut microbiome in the caterpillar *Manduca sexta*

**DOI:** 10.1101/2020.04.09.034165

**Authors:** Laura E. McMillan, Shelley A. Adamo

## Abstract

For many animals the gut microbiome plays an essential role in immunity and digestion. However, certain animals, such as the caterpillar *Manduca sexta*, do not have a resident gut microbiome. Although these animals do have bacteria that pass through their gut from their natural environment, the absence of such bacteria does not reduce growth or survival. We hypothesized that *Manduca sexta* would sterilize their gut as a protective measure against secondary infection when faced with a gut infection, or exposure to heat-killed bacteria in the blood (hemolymph). However, we found that gut sterilization did not occur during either type of immune challenge, i.e. bacterial numbers did not decrease. By examing the pattern of immune-related gene expression, gut pH, live bacterial counts, and weight change (as a measure of sickness behaviour), we found evidence for physiological trade-offs between between regulating the microbiome and defending against systemic infections. Caterpillars exposed to both gut pathogens and a systemic immune challenge had higher numbers of bacteria in their gut than caterpillars exposed to a single challenge. Following a principal component analysis, we found that the response patterns following an oral challenge, systemic challenge or dual challenge were unique. Our results suggest that the immune response for each challenge resulted in a different configuration of the immunophysiological network. We hypothesize that these different configurations represent different resolutions of physiological trade-offs based on the immune responses needed to best protect against the present immune challenges.

**SUMMARY STATEMENT:** This paper investigates the strategies that animals may use to regulate their microbiome during infection.

## INTRODUCTION

The gut microbiome is vital for many animals, both vertebrate (Montalban-Arques et al., 2015; Youngblut et al., 2019) and invertebrate (Fraune and Bosch, 2010; Weiss and Aksoy, 2011). The gut microbiome in insects is involved in: immune priming (Contreras-Garduno et al., 2016), supplying nutrients lacking in the diet (Brune, 2014), protecting hosts against parasites (Weiss and Aksoy, 2011) and harsh environmental conditions (Ferguson et al., 2018). However, these very same bacteria can become pathogenic if they are not regulated (e.g. if they reproduce without control and/or are allowed to cross into the blood (Buchon et al., 2014). Additionally, it has been shown in some insects (e.g. the greater wax moth -*Galleria mellonella*) that the presence of bacteria in the gut results in an increase of baseline AMPs which in turns has its own costs (Krams et al., 2017). During a systemic illness (i.e. an infection in the blood/hemolymph), the ability to control the gut microbiome wanes (Krieg, 1987). This decline in control increases the risk of a secondary infection via the gut (Krieg, 1987). Given this risk, how should animals deal with their gut microbiome during a systemic illness? The answer will depend, in part, on the role of the gut microbiome in a given species.

Although the gut microbiome is essential for survival in some species (e.g. termites, (Brune, 2014), in others it appears to be optional. For example, in the caterpillar stage of *Manduca sexta, Danaus chrysippus* and *Ariadne merione* destroying the gut microbiome with antibiotics has no effect on growth or development (Hammer et al., 2017; Phalnikar et al., 2019). In *Drosophila*, too, flies can grow without a microbiota, but only if provided with a rich food source (Buchon et al., 2014). However, even when they are fed a natural, relatively low quality, food source (i.e. *Datura wrightii*, (Ojeda-Avila et al., 2003) growth and survival is normal in *M. sexta* without a microbiome (Hammer et al., 2017). These results suggest that larval *M. sexta* do not require a gut microbiome.

Although gut bacteria do not appear to be necessary in *M. sexta*, they can still pose a threat. The gut is the main route of infection for most insects (Buchon et al., 2014; Vallet-Gely et al., 2008). Reducing the gut microbiome during gut infections may be important for survival. During severe immune responses within the gut, immunopathological damage can result in injuring tissues, as well as the delamination of enterocytes (Buchon et al., 2014). Damage to the gut increases the ability of gut bacteria to cross over into the hemocoel and cause lethal sepsis (Steinhaus, 1959).

Another potential reason to remove the microbiome is that having to regulate it requires immune resources in insects (Zhai et al., 2018) These costs could be problematic when *M. sexta* is facing multiple stressors simultaneously. Facing a combination of predators, toxins, or pathogens leads to a significantly increased likelihood of mortality than if each challenge was faced alone(McMillan et al., 2018). The increased mortality is due, in part, to physiological trade-offs between different defense systems (e.g. the immune system and stress response (Adamo et al., 2017) created by limited molecular resources (Adamo et al., 2016b; Adamo et al., 2017; McMillan et al., 2018). Therefore, in this species, it may benefit caterpillars to destroy their microbiome during a systemic infection to reduce the cost of maintaining it and reduce the risk of secondary infection. Sterilizing the gut is not without precedent in this species. *M. sexta* caterpillars sterilize their gut prior to metamorphosis (Russell and Dunn, 1996). During metamorphosis the gut is compromised, raising the real risk of attack by bacteria within the caterpillars system (Russell and Dunn, 1996). *M. sexta* sterilize their gut during metamorphosis by secreting large amounts of antimicrobial peptides and proteins into the gut lumen (Dunn et al., 1994; Russell and Dunn, 1996; Russell and Dunn, 1991). AMPs, such as lysozyme, accumulate in the gut epithelium. When the epithelium starts to slough off, the AMPs are released into the lumen, destroying the caterpillar’s microbiome (Russell and Dunn, 1996).

In this study, we examine whether a systemic infection leads to a loss of the transient microbiome, by measuring the bacterial content in the frass (fecal pellets) after a systemic immune challenge. We test whether systemic infection induces an upregulation of AMP gene expression in the midgut, similar to that observed during metamorphosis. We also test whether gut sterilization may be aided by an increase in gut pH. *M. sexta* typically have a highly alkaline gut with a pH ranging from 10-11 (Dow, 1992). An increase in pH could aid in reducing the number of bacteria in the gut by making the surrounding environment inhospitable.

Despite the potential advantages of removing the transient microbiome during a systemic immune challenge, there are also disadvantages. Sterilizing the gut is likely to be resource-intensive. Activating a systemic immune response requires resources, and has been found to be energetically costly in insects and other animals (Ardia et al., 2012; Bajgar et al., 2015; Lochmiller and Deerenberg, 2000), leading to physiological trade-offs (Krams et al., 2017; McMillan et al., 2018; Sheldon and Verhulst, 1996). Therefore, we also examine a second hypothesis that systemic immune activation will induce physiological trade-offs with microbiome regulation, leading to a reduction in midgut AMP gene expression and a reduction in gut alkalinity. Such trade-offs would be expected to produce an increase or maintenance of the number of bacteria in the gut during systemic immune activation, despite the risks.

A third possibility is that resource limitation may produce a reconfiguration of physiological networks as opposed to straightforward trade-offs. In some situations, *M. sexta* larvae adopt an alternate network configuration when faced with dual challenges (Adamo et al., 2016a; Adamo et al., 2017; McMillan et al., 2018). These alternative network strategies take the pressure off of molecular pinch points and increase survival (Adamo et al., 2016b; Adamo et al., 2017; McMillan et al., 2018). One possible alternate strategy in terms of microbiome regulation during an immune challenge would be to increase mechanisms of infection tolerance within the gut. Infection tolerance increases the ability of organisms to avoid damage due to pathogens, but it does not result in a reduction in pathogen load (Ayres and Schneider, 2009). Mechanisms of infection tolerance are less well studied than those of resistance, but include antioxidants and detoxification pathways (Soares et al., 2014; Soares et al., 2017). Reducing the chance of microbe-induced gut damage may help prevent gut bacteria from reaching the hemocoel. We examine whether systemically challenged caterpillars increase the expression of genes such as Gluthatione-S-Transferase (GST1). GST1 helps to detoxify compounds such as bacterial lipids (Snyder et al., 1995).

A fourth possible response is that *M. sexta* may use the microbiome to help it survive a systemic infection. Hammer et al.’s study (2017) showed that *M. sexta* grew normally without their gut microbiome, but this study did not assess whether the animals had a normal immune response. In Drosophila, there is evidence that the microbiota can participate in protecting against pathogens (Buchon et al., 2014). If *M. sexta* have similar mechanisms, than caterpillars may attempt to retain their microbiome during illness. In that case, during a systemic infection, the number of gut microbes should stay the same, and possibly even increase. We would not expect an increase in immune gene expression in the midgut.

To examine the likelihood of these 4 hypotheses, we exposed caterpillars to one of 5 treatments. Two of the groups underwent a single challenge, either oral inoculation with live bacteria (increasing the number and type of organisms in the microbiome) or injection of heat-killed pathogens (activating a systemic immune response). The other two treatment groups received dual challenges of oral inoculation of live bacteria combined with a sterile wound, or oral inoculation of live bacteria combined with an injection of heat-killed pathogens. The final group was an unmanipulated control. We measured gene expression in the midgut and fat body, as well as gut pH and microbial load.

Although *M. sexta* is unusual in not requiring a gut microbiome, the issue of how to deal with the gut microbiome during infection is a dilemma for all animals. This study helps us examine the costs and benefits of different microbiome control strategies in an animal that has a simpler microbiome than most other animals.

## MATERIALS AND METHODS

### ANIMALS

All studies were performed on 5^th^ instar larvae of *M. sexta* obtained from our colony. The colony was derived from eggs supplied by Great Lakes Hornworms (MI, USA), and was maintained as previously described (Adamo et al., 2016b). Trial caterpillars were weighed after their molt to the last larval instar (5^th^ instar- Day 0). Caterpillars were allotted into groups by weight, such that there were no initial weight differences across groups. Studies were approved by the University Committee on Laboratory Animals (Dalhousie University; I-11-025) and were in accordance with the Canadian Council on Animal Care.

### BACTERIAL INOCULATION

5th instar-Day 0 caterpillars were weighed and sorted into five different treatment groups at two timepoints (4 and 24 hours): 1) Control 2) Oral inoculation 3) Oral inoculation + Wounding challenge (control for systemic inoculation) 4) Systemic inoculation 5) Oral + Systemic inoculation.

Bacteria for oral inoculation, *Micrococcus luteus* (Microkwik culture, Carolina Biologica, Burlington, NC, USA) and *Enterobacter aerogenes* (Microkwik culture, Carolina Biologica) were chosen based on the types of mildly pathogenic bacteria previously found in field populations of lepidopterans (Hammer et al., 2017; Paniagua Voirol et al., 2018). Bacteria for systemic inoculation were a mixture of heat-killed *Serratia.marcescens* (Gram-negative bacterium, Microkwik culture, Carolina Biological, 1/10 LD50), *Bacillus cereus* (Gram-positive bacterium, Microkwik culture, Carolina Biological, 1/10 LD50) and *Beauveria bassiana* (strain GHA, fungus, 1/10 LD50, BotaniGard 22WP; Laverlam, Butte, MT, USA).

All 5^th^ instar-Day 0 caterpillars were given a 2.5 mm^3^ cube of high nutrition diet that had been dyed green with food colouring (ClubHouse, London, ON, Canada). For all groups except the Control and Systemic inoculation groups, the cube had been injected with a solution containing ∼10^6^ bacterial cells (50:50 *M. luteus: E. aerogenes)*. Bacteria numbers were estimated by optical density 600 nm and confirmed by plating on high nutrition agar. The remaining groups received a green food cube with the same dimensions but without the bacterial solution. Each caterpillar was given one hour to fully consume the cube. Any caterpillars who failed to consume the full cube were excluded from the study. Following this, all groups were given uncoloured high nutrition diet *ad libitum* for the following 23 hours. On 5^th^ instar-Day 1, all caterpillars had their food removed for 1 hour prior to manipulation and were weighed. The feeding protocol was then repeated. After the consumption of the green food cube was completed, Control and Oral inoculation were placed on uncoloured high nutrition diet for 4 or 24 hours depending on the group. Oral inoculation + wounding challenge caterpillars were given a sterile poke between the 6^th^ and 7^th^ abdominal segments before being placed on uncoloured high nutrition diet. Systemic inoculation and Systemic + Oral inoculation groups were given an injection of 20 µL of heat-killed triple mix (See above) between the 6^th^ and 7^th^ abdominal segments before being placed on uncoloured high nutrition diet.

### ORAL INOCULATION AND BACTERIAL COLONIES

In a pilot study we found that oral inoculation of bacteria significantly increased the culturable bacteria in both midgut contents (N=10) and frass (N=10) compared to controls (N=10 per contrast). Using fecal samples as a proxy method for non-invasive gut microbiome has been previously established in lepidoptera as well as *Drosophila* (Fink et al., 2013; Schwarz et al., 2018).

Frass pellets from an Orally inoculated group as well as Control group were collected, suspended in phosphate buffered saline and plated on nutrient agar. The gut transit time of the green coloured food bolus was found to be ∼2 hours regardless of the presence of inoculated bacteria which is consistent with previous studies (McMillan et al., 2018). Frass samples from all groups were collected and plated in duplicate at the following timepoints post-ingestion: 1 hour, 2 hours, 6 hours, 24 hours, and 72 hours. Agar plates were kept isolated at 23°C and colonies were counted 48 hours post plating.

### TISSUE SAMPLING AND RNA EXTRACTION

Caterpillars were sacrificed and dissected for midgut and fatbody tissue collection either 4 hours, or 24 hours post 5^th^ instar-day 2 manipulation. Caterpillars were chilled to induce a cold coma, and decapitated. The midgut was removed from the caterpillar, the midgut contents and peritrophic membrane were placed in a microcentrifuge tube. The midgut tissue was washed three times using ice-cold PBS before being placed in RNA*later* solution (Invitrogen, Carlsbad, CA, USA). Fatbody was extracted from the 7^th^ abdominal segment and placed in RNA*later* solution. Samples were kept in −80°C until being processed.

RNA extraction was performed using the RNeasy lipid tissue mini kit (Qiagen, Hilden, Germany). All steps adhered to the manufacturer’s instructions and included a DNase 1 treatment (RNase-Free DNaset, Qiagen) step to remove genomic DNA contamination. The integrity of total RNA samples was assessed using denaturing bleach gel electrophoresis (Aranda et al., 2012). The purity and concentration of extracted total RNA was determined using an Epoch Microplate Spectrophotometer (BioTek, Winooski, VT, USA) as well as a Qubit Fluorometer (Q32857, Invitrogen, CA, USA). Only samples with an A260/280 ratio greater than 1.8 were used. cDNA was synthesized using iScript (Bio-Rad, Hercules, CA, USA) and samples were stored at −20°C. Primers were purchased from integrated DNA technologies (http://www.idtdna.com/site) and stored at −20°C at a working stock of 10 µmol l−1. For primer sequences and efficiencies please see Table S1.

Prior to qPCR, each sample was diluted to a set concentration of 100 ng µl−1 using the Qubit Fluorometer (Q32857, Invitrogen). For each biological sample and gene, a 16 µl reaction mixture was prepared containing 4 µl of sample cDNA, 10 µl SsoAdvanced Universal SYBR Green Supermix (Bio-Rad), 0.6 µl of forward primer (10 µmol l−1), 0.6 µl of reverse primer (10 µmol l−1) and 0.8 µl RNase-free ddH2O. Reactions were performed in 96-well plates with a CFX96 real-time system (Bio-Rad). The reaction proceeded as follows: initial denaturation (95°C: 3 min), followed by 45 cycles of denaturation (95°C: 10 s), annealing and extension (60°C: 45 s). After the qPCR, a melt curve analysis was run to assess the specificity of the qPCR product. Quantitative cycle (ΔΔCq) values for each sample and gene target were calculated in CFX Maestro (Bio-Rad).

For reference gene assessment, we selected the most stable of six candidate reference genes used in a previous study in *M. sexta* (Adamo et al., 2016): Rp17A, actin (MSA), ribosomal protein S3 (MsS3), ubiquitin, beta FTZ-F1 and glycerol-3-phosphate dehydrogenase (G3PDH). We used NormFinder for R (http://moma.dk/normfinder-software) to determine stable reference genes (Andersen et al., 2004) (i.e. Rp17A and ubiquitin), using the Cq values of five biological samples for each candidate reference gene, for each treatment. The qPCR efficiency (E) and correlation coefficient (R2) for primer sets were estimated from a standard curve generated with 10-fold dilutions of mixed cDNA samples and are given in Table 1.

**Table 1.** Effect of systemic and oral challenges on *Manduca sexta* caterpillars 4 hours post treatment.

|  | Control<br>(N=14) | Oral+Wounding<br>(N=14) | Oral<br>(N=14) | Systemic<br>(N=14) | Oral+Systemic<br>(N=14) | *F* | *P* |
| --- | --- | --- | --- | --- | --- | --- | --- |
| Δ Mass | 0.72 ± 0.23 | 0.68 ± 0.22 | 0.54 ± 0.27 | 0.37 ± 0.08 * | 0.26 ± 0.32 * | 9.28 | 5.0 x 10<sup>-6</sup> |
| Colonies | 19.35 ± 14.7 | 69.32 ± 72.7 * | 28.57 ± 9.01 | 45.57 ±<br>47.46 | 165.36 ± 42.28 * | 25.24 | 1.22 x 10<sup>-12</sup> |
| pH | 10.27 ± 0.12 | 10.04 ± 0.11 * | 10.22 ± 0.22 | 9.86 ± 0.31 * | 10.07 ± 0.22 * | 9.23 | 2.0 x 10<sup>-5</sup> |
| FB Attacin | 1.23 ± 0.83 | 3.74 ± 0.66 * | 1.11 ± 1.18 | 4.94 ± 0.24 * | 4.71 ± 0.32 * | 90.29 | 8.22 x 10<sup>-26</sup> |
| MG Attacin | 0.63 ± 1.10 | 1.13 ± 0.79 | 0.74 ± 0.98 | 2.30 ± 0.18 * | 2.23 ± 0.26 * | 15.70 | 4.86 x 10<sup>-9</sup> |
| FB Lysozyme | 0.34 ± 0.30 | 1.49 ± 0.52 * | 0.26 ± 0.60 | 1.85 ± 0.12 * | 1.70 ± 0.23 * | 53.12 | 8.38 x 10<sup>-20</sup> |
| MG Lysozyme | 0.59 ± 0.75 | 2.13 ± 0.65 * | 1.28 ± 0.90 * | 2.59 ± 0.38 * | 2.76 ± 0.37 * | 28.06 | 1.49 x 10<sup>-13</sup> |
| FB GST | -0.13 ± 0.15 | 0.22 ± 0.08 * | -0.20 ± 0.20 | -0.24 ± 0.30 | -0.36 ± 0.26 * | 14.46 | 1.70 x 10<sup>-8</sup> |
| MG GST | -0.16 ± 0.24 | -0.12 ± 0.35 | -0.21 ± 0.23 | -0.22 ± 0.27 | -0.45 ± 0.29 * | 3.0 | 0.025 |
| FB Transferrin | 0.23 ± 0.26 | 0.78 ± 0.40 * | -0.03 ± 0.48 | 0.49 ± 0.24 | 0.25 ± 0.45 | 9.12 | 7.0 x 10<sup>-6</sup> |
| MG Transferrin | 0.25 ± 0.29 | 0.08 ± 0.50 | 0.22 ± 0.34 | 0.17 ± 0.33 | 0.20 ± 0.29 | 0.47 | 0.76 |
| FB NOS | -0.75 ± 0.41 | -0.08 ± 0.58 * | 1.11 ± 1.18 * | 0.21 ± 0.29 * | 0.38 ± 0.50 * | 14.65 | 1.40 x 10<sup>-8</sup> |
| MG NOS | -0.23 ± 0.46 | -0.22 ± 0.29 | -0.32 ± 0.33 | -0.18 ± 0.19 | -0.08 ± 0.30 | 1.08 | 0.375 |

Univariate analysis, followed by Dunnett's *post hoc* tests. Significance adjusted using Benjamini-Hochberg procedure.
Values are means ± s.d. <\*> denotes significance from control. FB, fat body; MG, midgut tissue. Δ Mass given in g, Colonies
given in CFUs and ΔΔCq values are given for qPCR results.

### pH MEASUREMENTS

The food bolus removed from the midgut was vortexed briefly to homogenize the sample, then the pH was measured using an StMicro5 pH electrode (OHAUS, Parsippany, NJ, USA) attached to a Corning pH meter 430 (Corning, Corning, NY, USA). All pH measurements were taken within 5 minutes of removal from caterpillar.

### BACTERIAL COLONIES AND TREATMENT GROUPS

Caterpillars had their abdominal segments surface sterilized with 70% ethanol and were placed in a disinfected container. Frass was collected immediately post excretion for 1 hour prior to tissue collection. Therefore, contamination from the outside environment should be minimal. The frass pellets were suspended in PBS and plated in duplicate on nutrient agar. The plates were isolated and allowed to grow at 23°C. 48 hours post plating bacterial colony forming units were counted.

In a pilot, frass was sent away for 16S metagenomic sequencing in order to better determine bacterial diversity, but amount of bacterial DNA was below the detection limit for a single individual. Data per individual was required for multivariate analysis.

### STATISTICS

Data were analyzed using SPSS (ver. 25). Data met the assumptions for MANOVA and univariate tests. Data for mass, pH, and bacterial colonies were found to be normally distributed using a Shapiro-Wilk test. The qPCR data were analyzed using CFX Maestro 1.1 (BioRad) and the REST program (2009; http://rest.gene-quantification.info). ΔΔCq values were log transformed prior to statistical analysis. When multiple tests were performed on the same dataset, the alpha criterion was corrected (Benjamini and Hochberg, 1995). Sample sizes were determined based on effect sizes derived from pilot data or literature values.

We used a multivariate approach (principle components analysis, PCA) to examine the pattern of activity across the dependent variables as described in (Stahlschmidt and Adamo, 2015). All data were log transformed to ensure equal variance across groups (Levene’s homogeneity of variances). Bartlett’s test of sphericity was significant for both time points (p<0.01). The correlation matrix (see Supplementary Table 3, 5) showed no extreme multicollinearity. The Kaiser-Meyer-Olkin Measure (>0.6 for both time points) demonstrated sampling adequacy. The first three principle components extracted from the data (eigenvalues >1.0) explained more than 60% of the variance for both time points. The factor scores for PC1, PC2 and PC3 were used to form 3-D plots.

## RESULTS

### BACTERIAL GUT TRANSIT TIMES

Gut transit times were not altered by the presence of bacteria. The gut transit times of high nutrition artificial diet inoculated with ∼10^6^ bacterial cells (50:50 *M. luteus: E. aerogenes)* was 135.5 ± 23 minutes while for control animals it was 125.2 ± 23 minutes (independent samples t-test, n=20, p=0.33). The number of colony forming units in the frass was higher for caterpillars fed bacteria, and increased over time (Fig. 1, 1 way-ANOVA timepoint; F(4,95)=1052.86, p<0.0001), with the exception of the 72 Hour post inoculation group in which the CFUs dropped to the bacterial count found between the 2 and 6 Hour timepoint (Figure 1).

**Figure 1.**
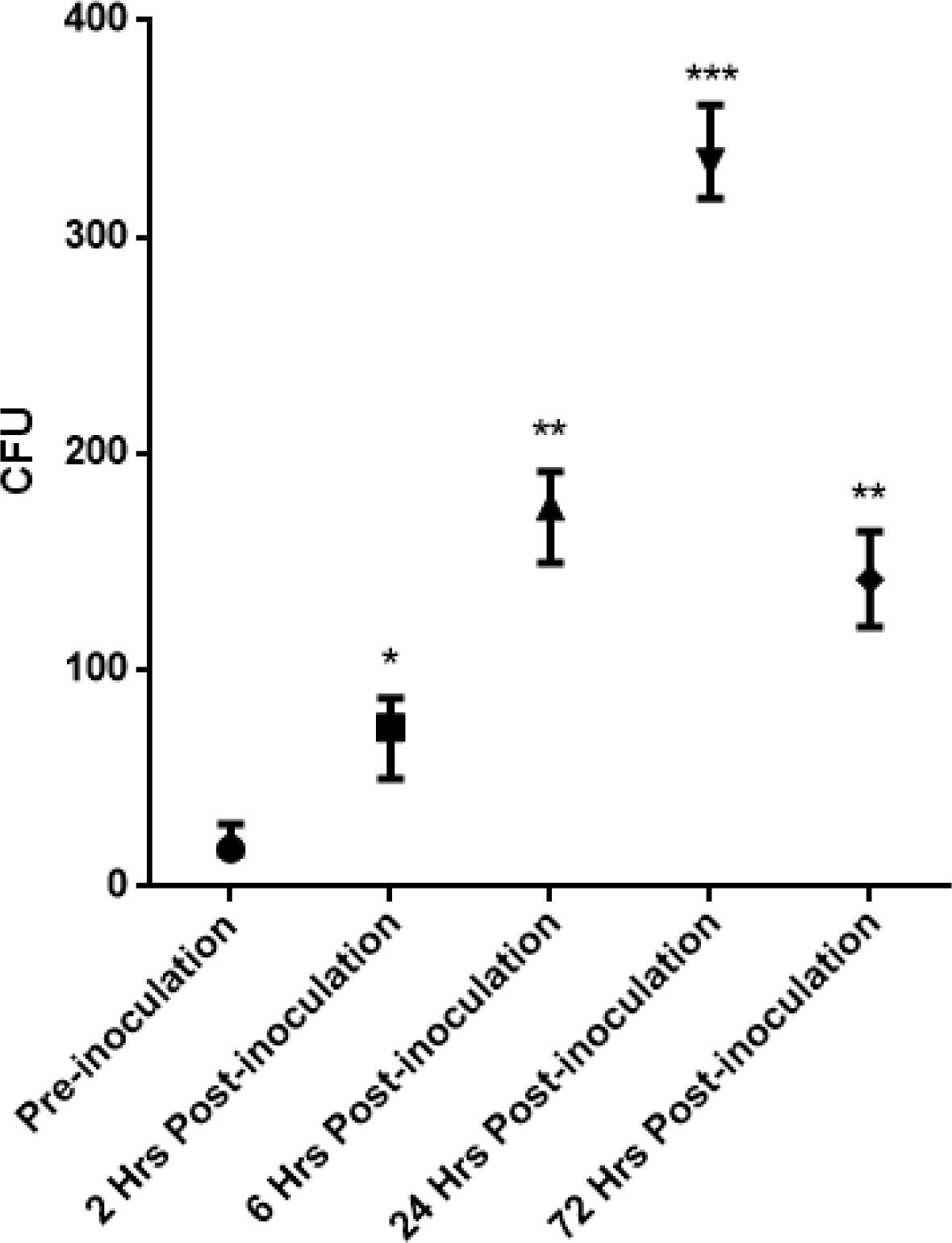
Bacteria gut transit times in *Manduca sexta.* Frass was collected and plated on high nutrient agar from *Manduca sexta* caterpillars prior to oral inoculation of bacteria and at several times points after. The number of colony forming units (CFUs) increased at each timepoint up to 24 hours post oral inoculation before decreasing again at 72 hours post oral inoculation For all groups n=20. The asterisk represent significant differences. Data points represent means and error bars represent the standard deviation.

The amount of CFUs present in the frass of Systemic+Oral challenge group was significantly higher level than that of control animals indicating that the bacteria travelling through the gut is thriving.

### FOUR HOUR TIMEPOINT

For each treatment group, change in mass (g), frass bacteria CFUs, gut pH, and ΔΔCq values for fat body and midgut gene expression were collected (Attacin, Lysozyme, GST1, iNOS, Transferrin). A MANOVA showed that there was a significant effect of treatment group (Wilks’ Lambda=0.002, F(13,56)=15.65, p < 0.0001). Individual univariate analysis were then conducted on the variables with p-values corrected using the Benjamini-Hochberg equation (Benjamini and Hochberg, 1995) (Table 1).

Change in mass (g) is representative of the intensity of illness-induced anorexia, and a sign of immune activation (Sullivan et al., 2016). At 4 hours post-manipulation, both Systemic and Oral+Systemically challenged *M. sexta* gained significantly less mass than control animals, indicating that at this timepoint both of these groups are exhibiting illness-induced anorexia. Interestingly, oral inoculation of bacteria (50:50 *M. luteus: E. aerogenes*) alone did not elicit this sickness behaviour until the 24 h timepoint (Tables 1, 2).

**Table 2.** Effect of systemic and oral challenges on *Manduca sexta* caterpillars 24 hours post treatment.

|  | Control<br>(N=14) | Oral+Wounding<br>(N=14) | Oral<br>(N=14) | Systemic<br>(N=14) | Oral+Systemic<br>(N=14) | *F* | *P* |
| --- | --- | --- | --- | --- | --- | --- | --- |
| Δ Mass | 1.83 ± 0.40 | 1.33 ± 0.57 * | 0.91 ± 0.50 * | 1.05 ± 0.32 * | 1.41 ± 0.53 | 8.00 | 2.60 x 10<sup>-5</sup> |
| Colonies | 40.43 ± 12.13 | 324.57 ± 64.47 * | 277.50 ± 69.72<br>* | 76.86 ±<br>28.96 | 349.43 ± 50.04 * | 116.89 | 6.08 x 10<sup>-29</sup> |
| pH | 10.30 ± 0.13 | 9.99 ± 0.40 * | 10.15 ± 0.10 | 9.84 ± 0.28 * | 10.41 ± 0.12 | 13.04 | 7.5 x 10<sup>-8</sup> |
| FB Attacin | -0.01 ± 0.55 | 1.17 ± 0.30 * | 0.79 ± 0.37 * | 2.33 ± 0.40 * | 2.40 ± 0.17 * | 103.78 | 1.73 x 10<sup>-27</sup> |
| MG Attacin | 0.25 ± 0.45 | 0.79 ± 0.57 | 0.93 ± 0.84 * | 1.74 ± 0.56 * | 1.60 ± 0.40 * | 15.25 | 7.57 x 10<sup>-9</sup> |
| FB Lysozyme | 0.70 ± 0.31 | 1.24 ± 0.24 * | 0.81 ± 0.31 | 1.57 ± 0.28 * | 1.81 ± 0.26 * | 40.45 | 5.54 x 10<sup>-17</sup> |
| MG Lysozyme | 0.65 ± 0.35 | 1.38 ± 0.60 * | 1.07 ± 0.66 | 1.21 ± 0.40 * | 1.39 ± 0.38 * | 5.37 | 0.001 |
| FB GST | -0.10 ± 0.20 | -0.31 ± 0.09 * | -0.53 ± 0.18 * | -0.35 ± 0.18 * | -0.40 ± 0.16 * | 12.28 | 1.70 x 10<sup>-7</sup> |
| MG GST | -0.02 ± 0.27 | 0.08 ± 0.19 | -0.31 ± 0.22 * | -0.18 ± 0.29 | -0.17 ± 0.20 | 5.85 | 4.39 x 10<sup>-4</sup> |
| FB Transferrin | -0.29 ± 0.19 | -0.10 ± 0.25 | -0.09 ± 0.25 | -0.03 ± 0.23 * | 0.21 ± 0.23 * | 8.43 | 1.50 x 10<sup>-5</sup> |
| MG Transferrin | -0.92 ± 0.62 | -0.67 ± 0.36 | -0.86 ± 0.33 | -0.87 ± 0.35 | -0.25 ± 0.62 * | 4.80 | 0.002 |
| FB NOS | -0.06 ± 0.20 | 0.14 ± 0.26 | 0.18 ± 0.23 * | 0.34 ± 0.33 * | 0.27 ± 0.19 * | 5.37 | 0.001 |
| MG NOS | -0.03 ± 0.27 | -0.13 ± 0.21 | -0.15 ± 0.33 | -0.26 ± 0.20 | -0.12 ± 0.27 | 1.39 | -0.246 |

Univariate analysis, followed by Dunnett's *post hoc* tests. Significance adjusted using Benjamini-Hochberg procedure.
Values are means ± s.d. <\*> denotes significance from control. FB, fat body; MG, midgut tissue. Δ Mass given in g, Colonies
given in CFUs and for qPCR ΔΔCq. For the qPCR data, values are normalized against the reference genes.

The gut pH of *M. sexta* is very basic, averaging at about 10-11 on the pH scale (Dow, 1992). Four hours post treatment, *M. sexta* in the Oral+Wounding challenge, Systemic challenge, and Oral+Systemic challenge groups all dropped their gut pH compared with controls, making their midgut significantly less basic than control animals. *M. sexta* that had been orally inoculated alone did not have this drop in midgut pH, but did not increase pH either (Table 1).

The frass that was collected and plated 4 hours post manipulation showed a significant increase in the amount of CFUs present in *M. sexta* in the Oral+Wounding challenge and Oral+Systemic challenge groups compared with controls.

Within the fat body, attacin and lysozyme gene expression were upregulated in the Oral+Wounding challenge, Systemic challenge, and Oral+Systemic challenge groups. Oral inoculation did not elicit the production of AMPs within the fat body at this timepoint (Table 1, Figure 2ab). Within the midgut tissue, attacin was upregulated in the Systemic challenge and the Oral+Systemic challenge groups. Lysozyme expression in the midgut was upregulated in all groups, relative to controls, at 4 hours post manipulation (Table 1, Figure 2cd).

**Figure 2.**
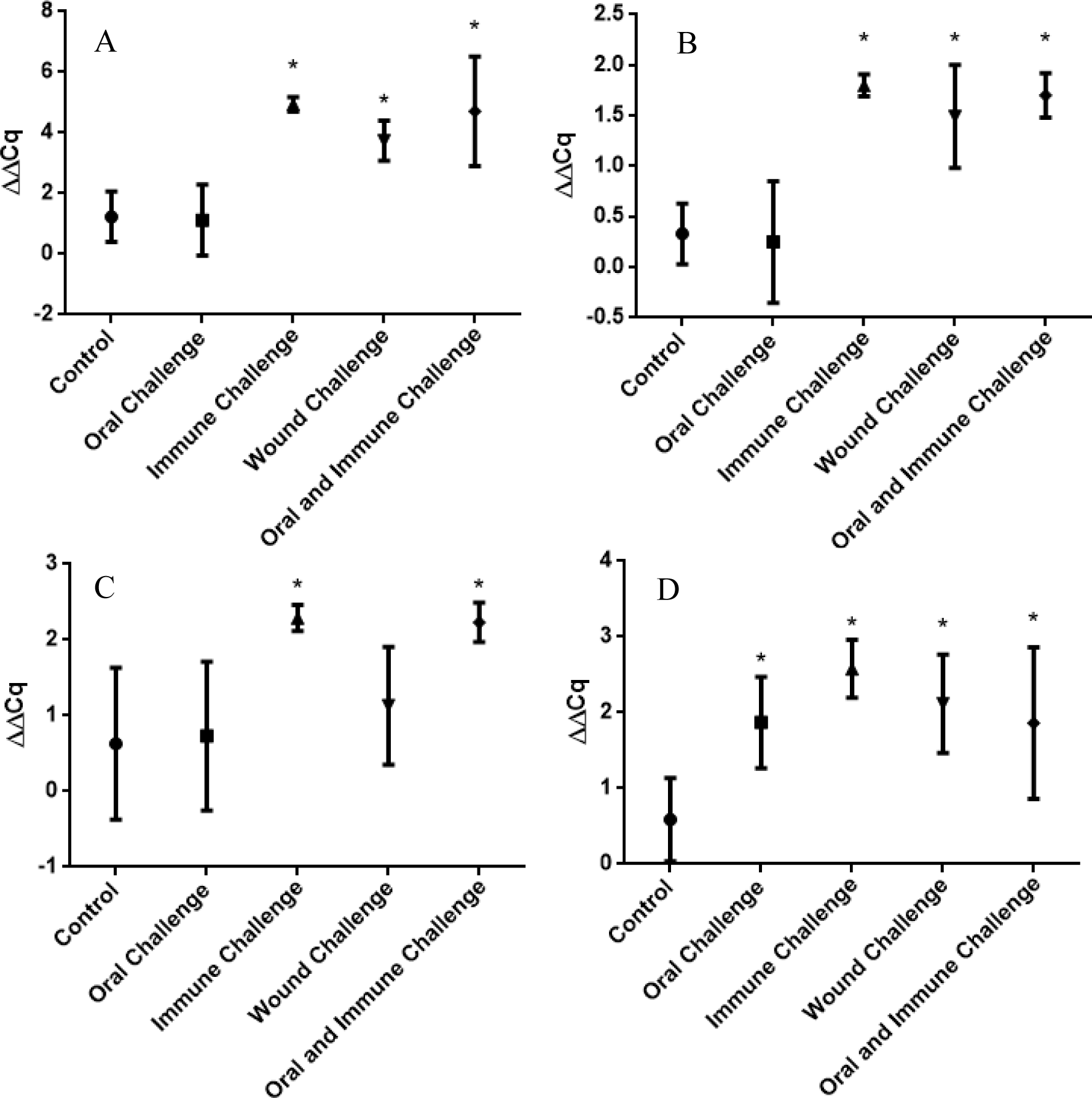
Antimicrobial peptide mRNA expression in fat body and midgut tissues at 4 hours post manipulation. Expression of the AMP attacin in the fat body (A) and midgut tissue (B). Expression of the AMP lysozyme in the fat body (C) and midgut tissue (D). Significance is denoted with an *. Data points denote the average and error bars represent standard deviation.

The transcription levels of the mRNA of GST1 showed interesting patterns of regulation, with the expression being upregulated in the Oral+Wounding challenge group at 4 hours post manipulation and being significantly downregulated in the Oral+Systemic challenge groups (Table1, Figure 3a). Within the midgut tissues the same downregulation can be seen within the Oral+Systemic challenge, but the upregulation seen in the Oral+Wounding challenge group was not present (Table 1, Figure 3b).

**Figure 3.**
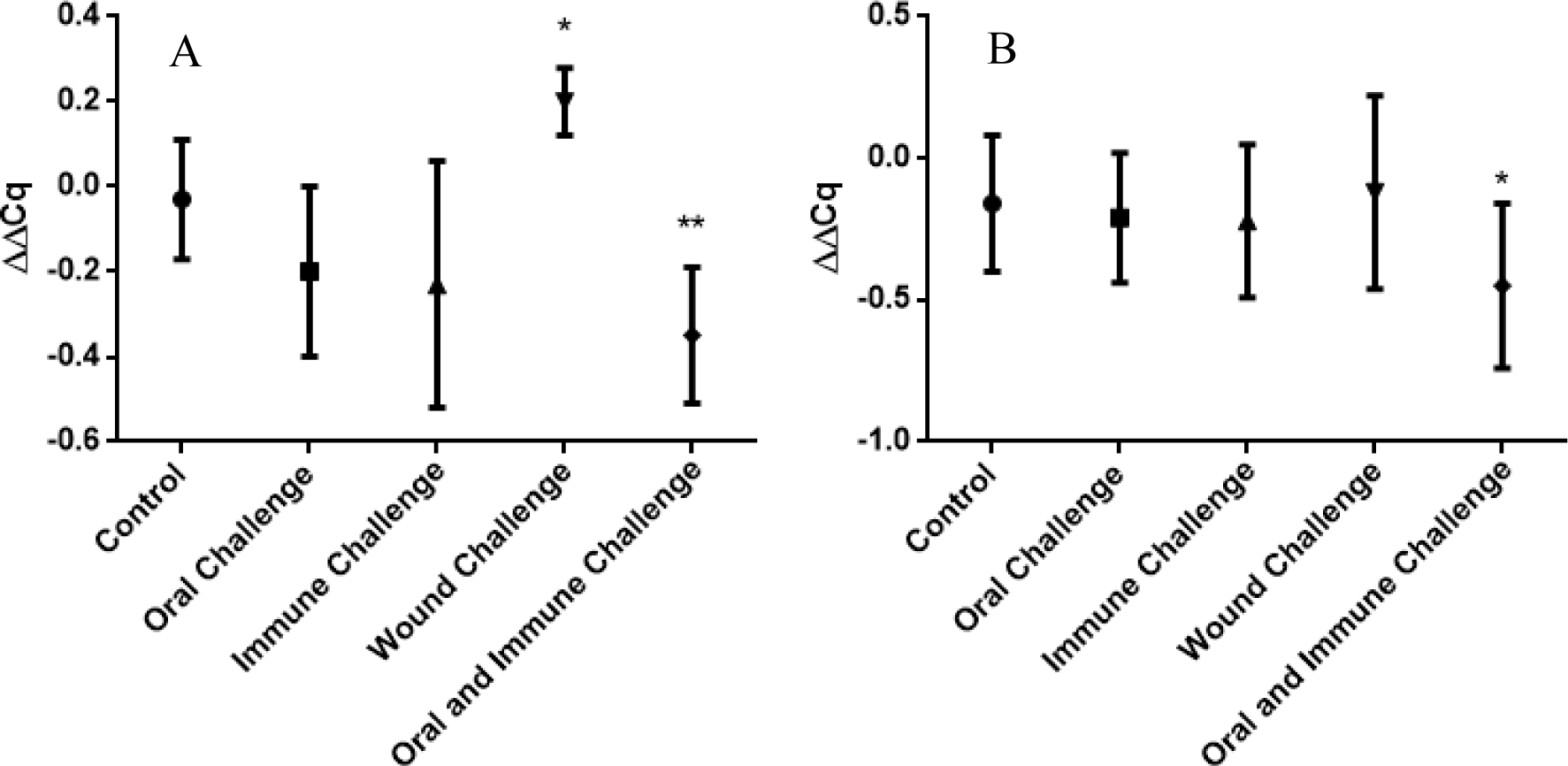
Glutathione-S-Transferase mRNA expression in the fat body and midgut tissues of *Manduca sexta* 4 hours post manipulation. Expression of GST1 in the fat body (A) and midgut tissues (B). Significant upregulation is denoted by *. Significant down regulation is denoted by **. Data points represent the mean and the error bars represent the standard deviation.

Inducible nitric oxide synthase (iNOS) was upregulated under all conditions in the fat body at this timepoint. However, in the midgut tissue, iNOS was not induced in any of the challenge groups (Table 1).

### TWENTY-FOUR HOUR TIMEPOINT

A MANOVA showed that there was a significant effect of treatment group (Wilks’ Lambda=0.002, F(13,56)=16.04, p < 0.0001). Individual univariate analysis were then conducted on the variables, with p-values corrected using the Benjamini-Hochberg equation (Benjamini and Hochberg, 1995) (Table 2).

At the twenty-four hour timepoint we expected to see a recovery from illness-induced anorexia. As predicted, the Oral+Systemic challenge group had recovered (Table 2). However, illness-induced anorexia appeared in the Oral+Wounding Challenge, and was still measurable in Oral challenge and Systemic challenge groups (Table 2).

The pH of the Oral+Wounding challenge and the Systemic challenge group remained less basic than the control group 24 hours post manipulation. However, the midgut pH of the Oral+Systemic challenged group was no longer different compared to controls (Table 2).

All groups that were orally inoculated with live bacteria had significantly higher CFUs in their frass than control animals (Table 2).

The mRNA expression for antimicrobial peptide attacin was upregulated in the fat body tissue in the Oral+Wounding challenge, Systemic Challenge and Oral+Systemic challenge groups. Unlike the 4 hour timepoint, at 24 hours post manipulation the Oral challenge group also showed increased expression of attacin (Table 2, Figure 4a). Lysozyme expression within the fat body remained upregulated for the Oral+Wounding challenge, Systemic challenge, and Oral+Systemic challenge groups (Table 2, Figure 4b). Within the midgut tissue, gene expression of attacin was upregulated in Oral challenge, Systemic challenge, and Oral+Systemic challenge groups. Lysozyme was upregulated in the Oral+Wounding challenge, Systemic challenge and Oral+Systemic challenge groups (Table 2, Figure 4cd).

**Figure 4.**
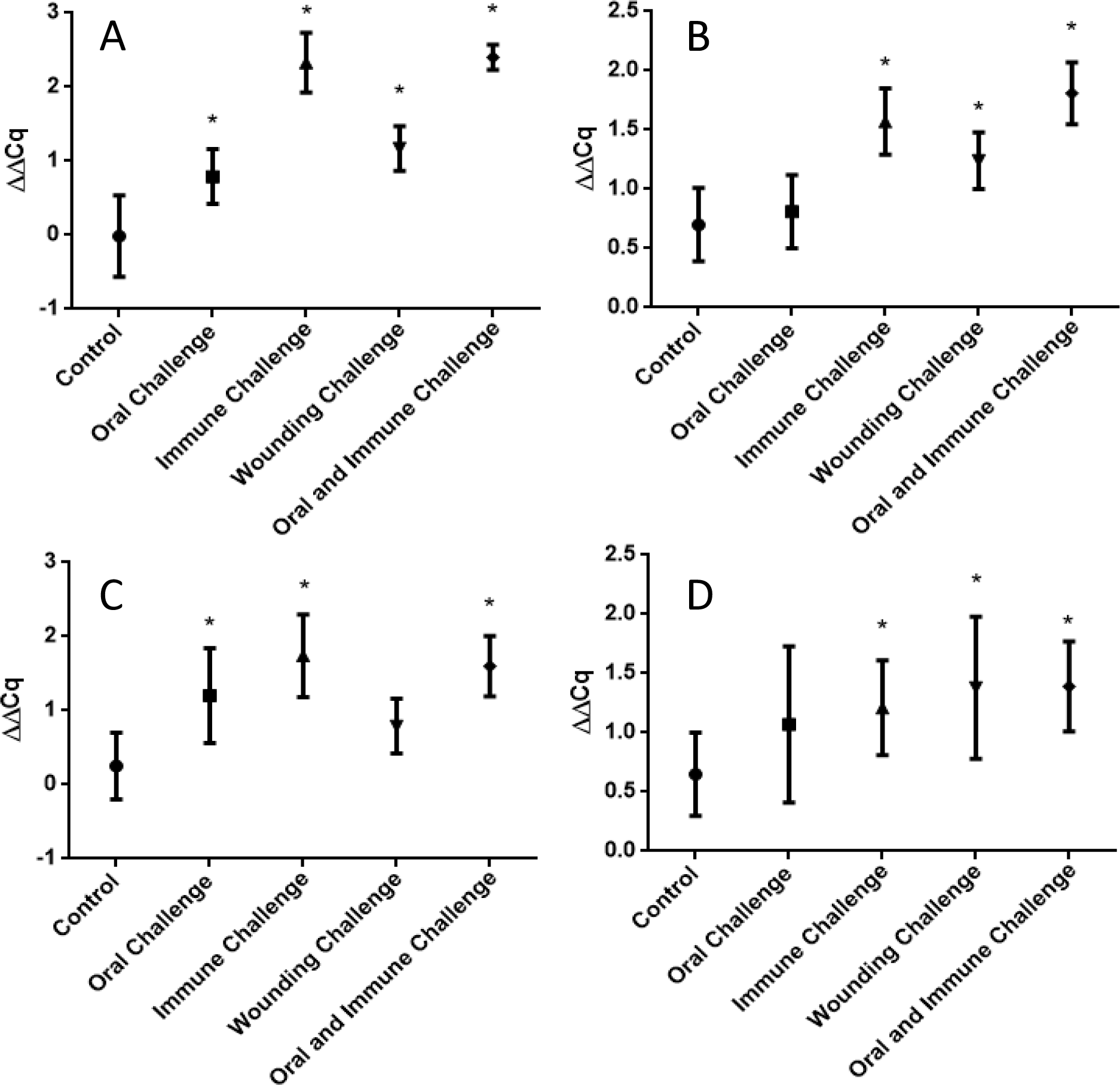
Antimicrobial peptide mRNA expression in fat body and midgut tissues 24 hours post manipulation. Expression of the AMP attacin in the fat body (A) and midgut tissue (B). Expression of the AMP lysozyme in the fat body (C) and midgut tissue (D). Significance is denoted with a *. Data points represent the mean and the error bars represent the standard deviation.

At 24 hours post manipulation, GST1 gene expression was downregulated in the fat body in each of the groups when compared to control animals (Table 2). This same trend was not seen in the midgut tissue, where at the 24 hour timepoint there was only downregulation in the Oral challenge group (Table 2).

Gene expression of iron-binding glycoprotein transferrin was upregulated in the fat body and midgut tissues of *M. sexta* that had undergone an Oral+Systemic challenge (Table 1).

The increased expression of nitric oxide synthase in the fat body tissue was still visible at 24 hours post manipulation in the Oral+Wounding challenge, Systemic challenge, and Oral+Systemic challenge groups. The expression of iNOS was no longer increased in the Oral challenge group when compared with controls (Table 2). In the midgut, there were no significant changes in iNOS expression at 24 h (Table 2).

### PRINCIPAL COMPONENTS ANALYSES

The complexity of the results led us to use a principal components analysis (PCA) to search for patterns in the data. A PCA at the four hour time point revealed that there are three main factors that explain the majority (62%) of the variation between our groups (Figure 5).

**Figure 5.**
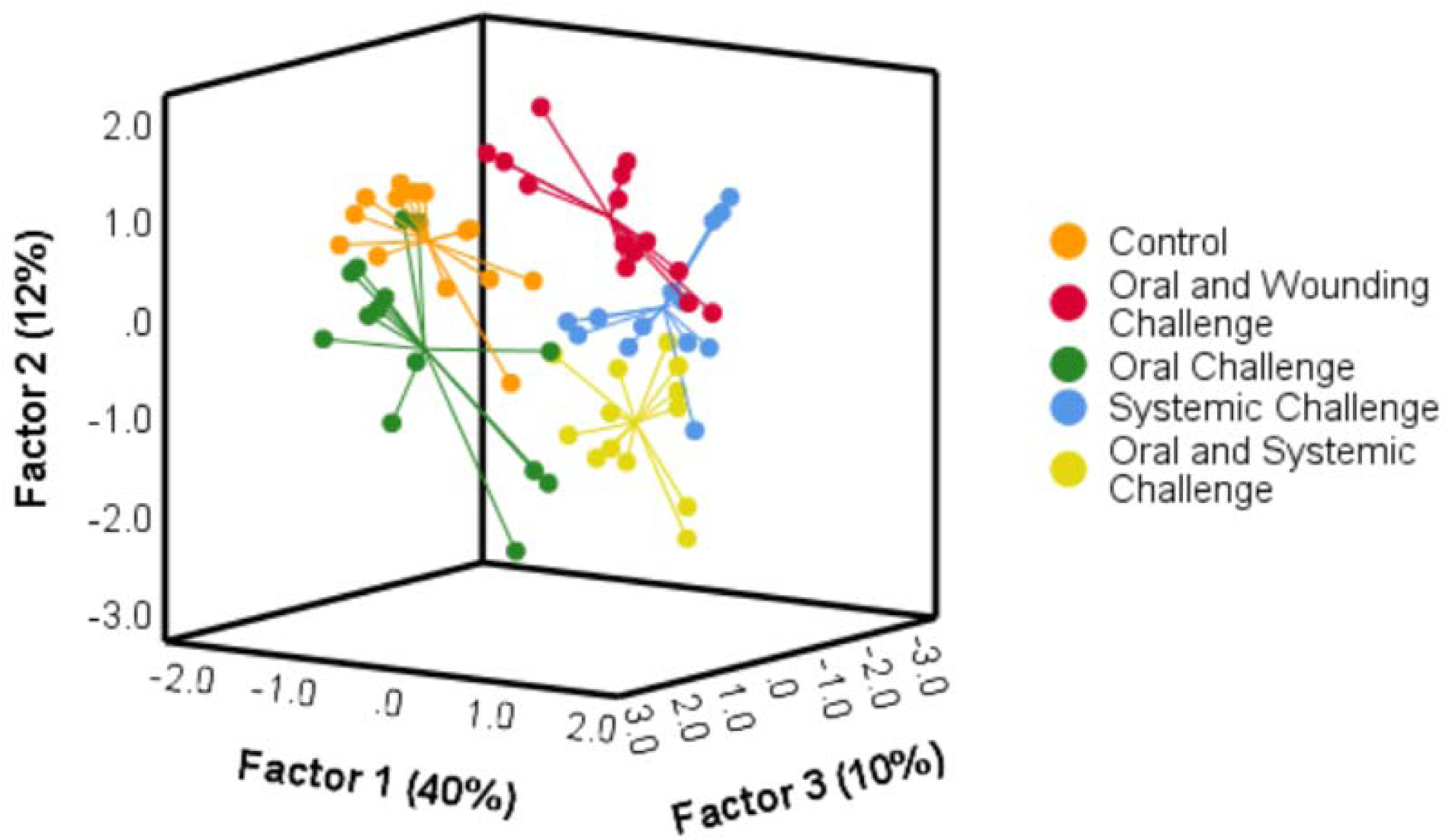
Principal components analysis of variables 4 hours post manipulation. Each point represents an individual *M. sexta.* Lines meet at the centroid of each group.

Each treatment resulted in a group with differing centroids within the PCA. Factor 1, which accounted for 40% of the variability between groups was highly correlated with AMPs present in both the fat body and the midgut tissues (Table S2, S3). It was negatively correlated with the expression of GST1 within the midgut tissue (Table S3).

At the 24 hour time point, the different treatments also resulted in separated centroids and groups with three main factors explaining the majority (60%) of the variation (Figure 6, Table S4). Factor 1, which account for 35% of the variability, for the 24 hour timepoint was also positively correlated with the AMP expression in both tissues and negative correlated with the expression of GST1 (Table S5). For both timepoints, the Oral+Systemic dual challenge is distinct from both Oral challenge and Systemic challenge groups (i.e. little (4 h) or no (24 h) overlap of datapoints within a cluster).

**Figure 6.**
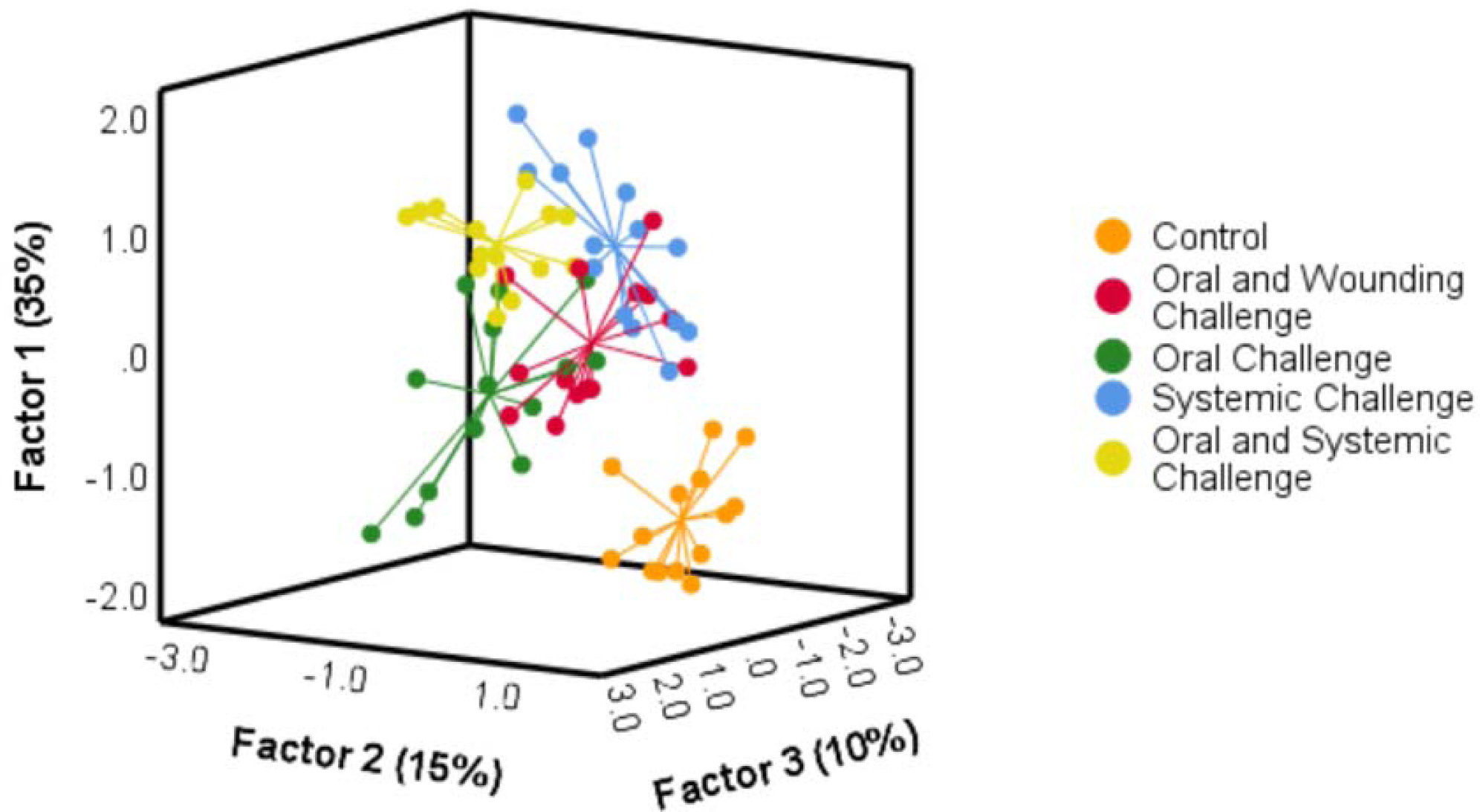
Principal components analysis of variables 24 hours post manipulation. Each point represents an individual *M. sexta.* Spikes meet at the centroid of each group.

## DISCUSSION

There was no evidence that *M. sexta* sterilizes its gut during either a Systemic, Oral or a dual immune challenge. Bacteria remained in the gut (as estimated from counting bacteria colony forming units in the frass) during both Systemic and Oral challenges. In fact, there was no evidence that the number of gut bacteria even declined during these challenges (Tables 1 and 2). A decline in gut bacteria numbers was expected in treatments with systemic immune challenges because the microbiome of *M. sexta* comes from its food (Hammer et al., 2017), and caterpillars eat less during an immune challenge (i.e. illness-induced anorexia). However, we saw no such decline. Although immune challenged caterpillars eat less, their gut transit time remains unchanged (McMillan et al., 2018). Therefore, the number of culturable bacteria in the frass was not artificially elevated by a decline in the movement of material through the gut. We also know that both the Oral and Systemic challenges produced robust immune responses in the caterpillars because of the expression of illness-induced anorexia and the increased expression of immune-related genes such as attacin (Tables 1 and 2). Therefore, the lack of gut sterilization was not due to a lack of an immune response. Finally, the immunological evidence that caterpillars attempted to sterilize their gut was equivocal. Although challenged caterpillars did demonstrate an increase in AMP production in both midgut and fat body, there was no increase in gut alkalinity (i.e. increase in gut pH). Instead of an increase in gut alkalinity, the Systemic challenge, Oral+Wounding challenge, and Systemic+Oral challenge groups dropped their gut pH compared to controls (Table 1,2). A less alkaline gut would make the gut environment more hospitable to bacteria such as *Enterobacter aerogenes* and *Micrococcus luteus* that function optimally at a more neutral pH (Kung and Wang, 1977; Tanisho, 1998). However, the decline in pH was modest (approximately 0.4 pH units). Whether this change would significantly enhance bacterial survival is uncertain. These results show that gut sterilization is not a host defense mechanism for pathogen threats in *M. sexta*, at least not for the immune challenges used in this study.

During severe sepsis, *M. sexta* produce watery feces, suggesting that they are attempting to flush their gut of pathogens (Dunn et al., 1994). We have only observed this in *M. sexta* close to death (pers. obs.). It is possible that removing bacteria from the gut remains an option for *M. sexta*, when all other defenses have failed.

Our data are more consistent with the three other hypotheses we explored. The first alternative hypothesis suggested that gut sterilization may not occur because of physiological trade-offs between systemic immune defense and the resources needed for sterilization. Trade-offs may even lead to a reduction in the mechanisms usually used to keep the microbiome in check, leading to an increase in the number of bacteria in the gut. However, in caterpillars given a Systemic immune challenge, we saw no evidence for an increase in the number of culturable bacteria in the gut compared with controls. However, it is possible that there was a change in the number of bacteria that do not grow on nutrient agar. Nevertheless, minimally we can conclude that there was no increase in some types of bacteria, suggesting that there was no decline in at least some aspects of immune defense in the gut. Nevertheless, there was evidence for a trade- off between systemic defense and bacterial numbers in the gut during a dual challenge. Bacterial numbers in the gut were higher during a dual challenge than an oral challenge. However, we did not see evidence of declines in the production of two important AMPs in the midgut (attacin and lysozyme) during a dual challenge, as might be expected if there were physiological trade-offs involving these molecules. In fact, gene expression of these AMPs was sometimes higher during the dual challenge compared with expression during a single challenge (Table 2, Figure 4). It is possible that trade-offs were occurring in unmeasured immune components.

The second alternative hypothesis suggests that *M. sexta* may maintain its transient gut microbiome because it confers benefits in terms of immunity (Fraune and Bosch, 2010). Although *Manduca sexta* do not appear to possess a resident gut microbiome, they do acquire a microbiome from their food and environment (Hammer et al., 2017), that could be beneficial even if it is transient. Our results showed that the transient gut microbiome is retained during both systemic and gut bacterial infections. Determining whether it is advantageous will require further study.

Caterpillars may also alter (i.e reconfigure) their immune response when faced with a dual challenge as opposed to a single challenge. Insects have been shown to reconfigure their immune response when an immune challenge co-occurs with environmental stressors such as food availability and predators (Adamo et al., 2008; Adamo et al., 2016b; Adamo et al., 2017). One possible reconfiguration would be a shift in emphasis from resistance mechanisms to infection tolerance mechanisms in the midgut. During a dual challenge, we predicted that the caterpillar would prioritize protection over resistance as resources for resistance mechanisms may be in short supply. Infection tolerance mechanisms may be less expensive than resistance mechanisms, and less damaging (Soares et al., 2014).

Glutathione-S-Transferases (GSTs) are important detoxification enzymes that perform a dual role. They are involved in detoxification of xenobiotics via catalyzing the conjugation of the toxin with glutathione (Eaton and Bammler, 1999; Snyder et al., 1995), as well as conferring protection against self-harm induced by oxidative stress pathways (Kim et al., 2011). However, Glutathione-S-Transferases are a large family of enzymes with 40 unique gene transcripts having been identified in *M. sexta* (Koenig et al., 2015). The transcripts of identified GST genes have been shown to cluster in a tissue specific manner, with some being found exclusively in the fatbody or midgut, well others such as GST1 being expressed in multiple tissues (Koenig et al., 2015; Snyder et al., 1995). GST1 was selected for this study, as it has been identified in the literature as being present in the midgut, and to show variable expression based on diet type, and presence of bacteria (Snyder et al., 1995). We predicted an increase in GST-1 production in the midgut during a dual challenge, because such an increase should enhance protection against bacterial toxins (Snyder et al., 1995). However, at the 4 h time point we saw a steep reduction in GST-1 expression in both fat body and midgut. Midgut GST-1 expression had recovered by 24 h. This pattern is not found in all insects. Cabbage loopers (*Trichoplusia ni*) show no change in GST-1 expression in the midgut after being fed bacteria, although a decline is found in other tissues (Freitak et al., 2009). It is difficult to make a definitive conclusion because we did not assess the activity of all GST genes active in the midgut. However, the ability of an immune challenge to induce a steep decline in a GST gene (i.e. GST-1) that plays a major role in detoxification (Snyder et al., 1995) may point to a physiological trade-off between detoxification and immune defense (McMillan et al., 2018).

Transferrin is a protein that sequesters free iron as part of a tolerance strategy called nutritional immunity (Soares and Weiss, 2015), and is another potential mechanism of infection tolerance (Brummett et al., 2017). *M. sexta* shows increased expression of mRNA for transferrin in the fatbody 24 hours after a bacterial injection into the hemocoel (Brummett et al., 2017; He et al., 2015). Our results support these findings. We found an increase of transferrin in the fatbody 24 hours post treatment in both the Systemic challenge group as well as the Oral+Systemic challenge group. Interestingly, ingesting bacteria did not increase transferrin expression in the midgut, although a dual challenge did (Table 2). This finding may indicate that a shift towards tolerance mechanisms in the midgut may only be activated during a widespread or massive immune challenge, e.g. during a combined systemic and gut infection. It is also possible that ingesting a more virulent bacteria could have induced transferrin expression.

Although we found some evidence for an increase in tolerance mechanisms in the gut during immune challenges, we assessed only a small number of possible mechanisms. Unfortunately tolerance mechanisms are poorly characterized (Ayres and Schneider, 2012). In the future, measurements of damage (e.g. signs of oxidative stress, (Costantini, 2019) may provide the best estimate of the activity of tolerance mechanisms.

On the other hand, our results do suggest an increase in resistance mechanisms in the midgut. We observed an increase in gene expression for the AMPs attacin and lysozyme, even during a systemic immune challenge (i.e. when there was no increase in bacteria in the gut). It is possible that the midgut becomes recruited to support the systemic immune system; there is evidence that AMPs secreted by the midgut can end up in the hemolymph (Freitak et al., 2007).

Another resistance mechanism used extensively in the insect gut is the production of reactive molecules (e.g. ROS) to destroy pathogens. Buchon and Cherry (2014) showed that *Drosophila* increase the amount of ROS in their gut in response to natural microbiota by activating DUOX and iNOS. Similarly, Eleftherianos et al. (2009) demonstrated in *M. sexta* that when the pathogen (*Photorhabdus luminescens*) was ingested, increased gene expression of iNOS was restricted to the gut. On the other hand, iNOS was increased in the fat body when pathogens were injected into the hemocoel (Eleftherianos et al., 2009). However, in our study, we found that iNOS mRNA expression increased significantly in the fatbody after the Oral challenge, Systemic challenge, and Oral+Systemic challenge groups at both 4 hours and 24 hours post treatment. We did not, however, see an increase in the expression of iNOS transcripts in the midgut tissues at either time point. It is possible that these differences are due to our using a primer designed for a specific inducible nitric oxide synthase which would not capture the whole picture if other iNOS genes are involved. Moreover, we used bacteria that were mildly pathogenic as the Oral challenge, and we injected heat-killed, not live, bacteria into the hemocoel for the Systemic challenge. These differences may account for the differences in the expression pattern.

The principal components analyses at both timepoints were most strongly influenced by the amount of AMPs being produced in both tissues (Table S3, S5). The analysis demonstrates that the pattern of changes among all the dependent variables were distinct across the five treatments, leading to different centroids at both timepoints. This result supports the hypothesis that the immunophysiological response to a challenge depends on the nature and context of the challenge (e.g. Adamo, 2017). The immune system responds differently depending on the number of simultaneous immune challenges, for example (Fig. 4). It is not uncommon for animals to be exposed to multiple pathogens (e.g. insects, (Boucias and Pendland, 2012). Therefore, natural selection would favour animals that could produce an optimal response to concurrent gut and systemic pathogen challenges. Optimally responding to different types and numbers of pathogens requires partitioning scarce resources among different immune components and multiple physiological systems. In animals with microbiomes, their regulation during an infection is likely to vary depending on the number and type of concurrent immune challenges.

## ACKNOWLEDGEMENTS

We thank L. Puddicombe and B. Chiang for help with animal care. We thank D. Miller for help with experimental design.

## DATA AVAILABILITY

All datasets can be accessed via Mendeley Data

## AUTHORS’ CONTRIBUTIONS

LEM helped plan the study, carried out experiments, data analysis, and helped draft the manuscript; SAA conceived of the study, helped design the study and helped draft the manuscript. All authors gave final approval for publication.

---

**Figure S1.**
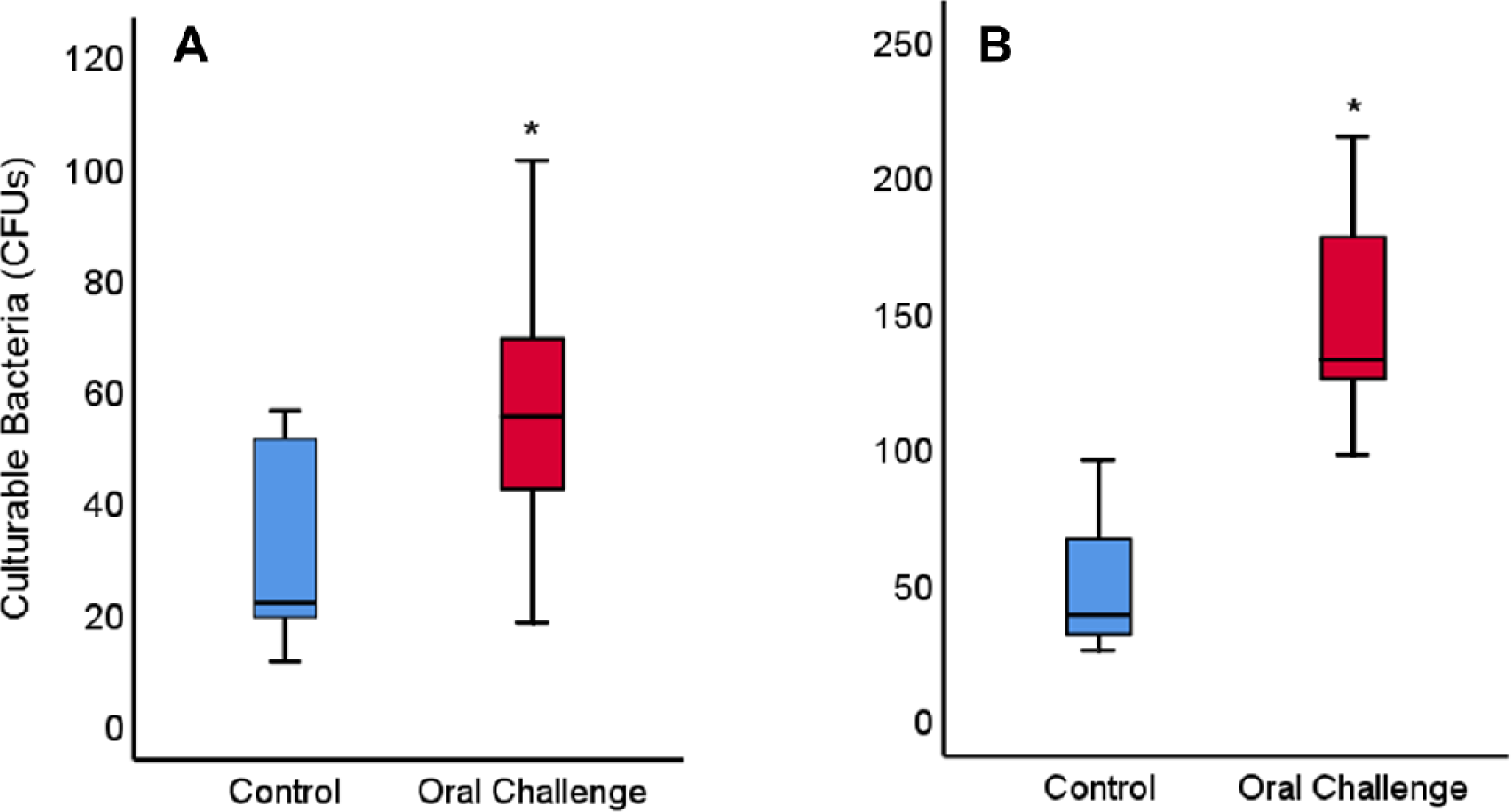
Cultrable bacteria from the midgut contents (A) and frass (B) of *M. sexta*, 4 hours post feeding. Oral challenge group were given a food cube that had been injected with a solution containing ∼10^6^ bacterial cells (50:50 *M. luteus: E. aerogenes)*. Independent samples T-test from midgut contents indicate a significantly greater amount of bacteria in the oral challenge (N=10, M=57.9, SD=24.7) group relative to the control group (N=10, M=29.1, SD=17.5); t(18)=-3.00, p=0.008. Independent samples T-test from plated frass similarly indicates a significantly greater amount of bacteria in oral challenge (N=10, M=147.5, SD=11.31) compared to control (N=10, M=47.0, SD=23.2); t(18)=-7.5, p<0.0001

**Table S1.**
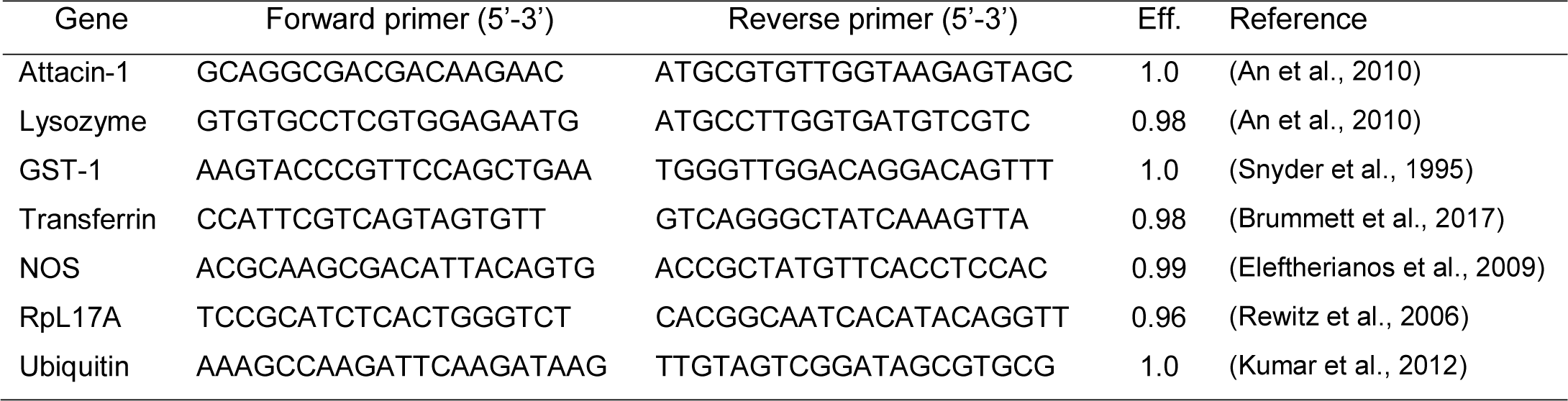
Forward and reverse primer sequences for target immune-related genes and reference genes.

**Table S2.** Principal component analysis for 4 hours post treatment

|  | Eigenvalues | % Variance |
| --- | --- | --- |
| Factor 1 | 5.0 | 39.68 |
| Factor 2 | 1.8 | 12.44 |
| Factor 3 | 1.3 | 9.88 |

**Table S3.**
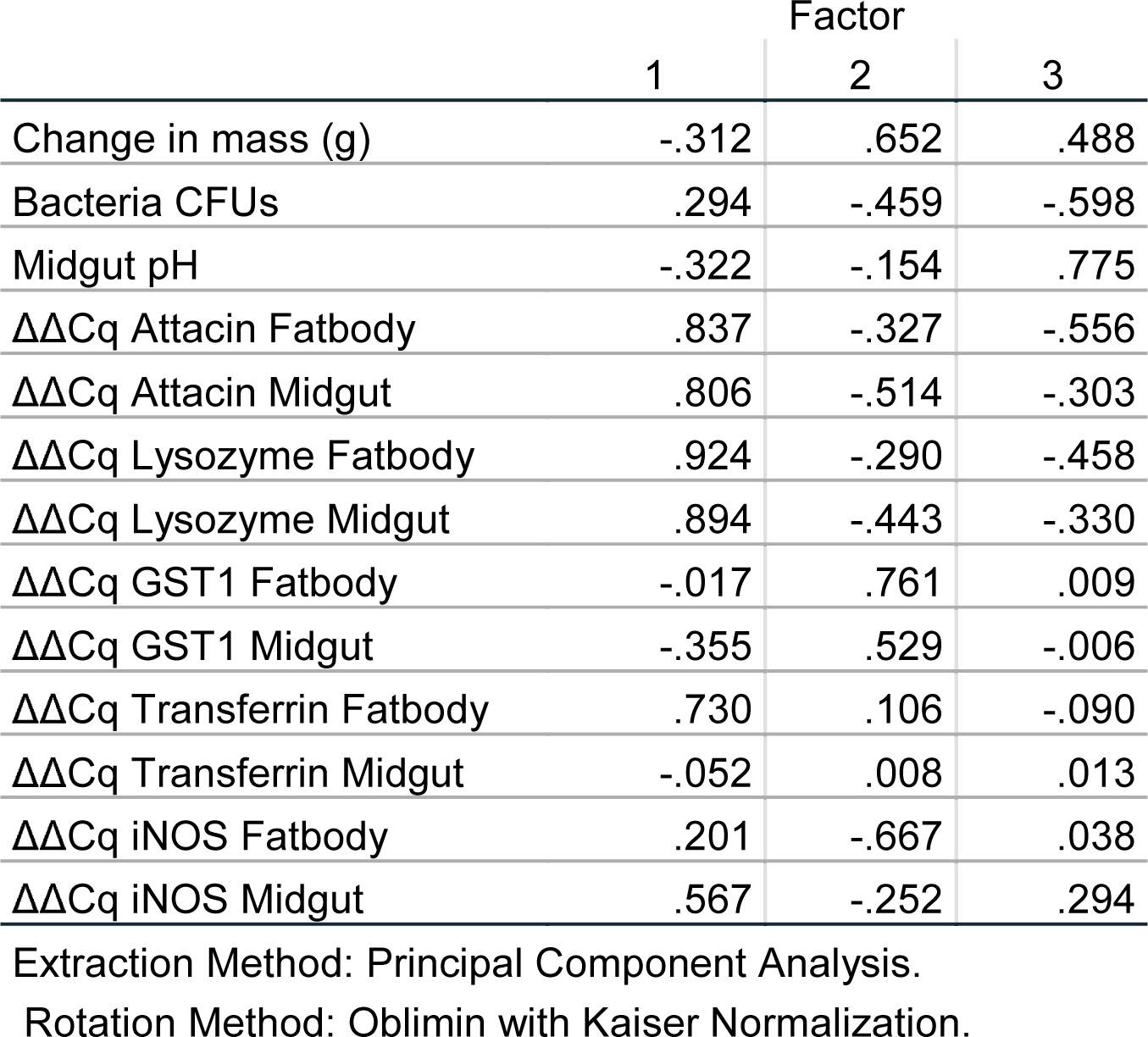
Correlation of traits for the three main factors of the 4 hour time point PCA

**Table S4.**
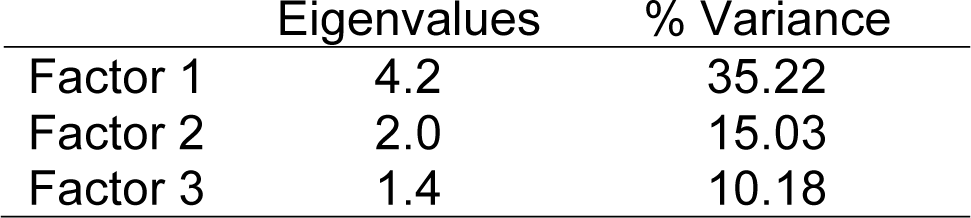
Principal component analysis for 24 hours post treatment

**Table S5.** Correlation of traits for the three main factors of the 24 hour time point PCA

|  | Factor |  |  |
| --- | --- | --- | --- |
|  | 1 | 2 | 3 |
| Change in mass (g) | -.117 | .793 | .124 |
| Midgut pH | -.212 | .199 | .694 |
| Bacteria CFUs | .476 | -.350 | .408 |
| $\Delta\Delta\text{Cq Attacin Fatbody}$ | .882 | -.302 | .224 |
| $\Delta\Delta\text{Cq Attacin Midgut}$ | .871 | -.194 | -.059 |
| $\Delta\Delta\text{Cq Lysozyme Fatbody}$ | .858 | -.092 | .314 |
| $\Delta\Delta\text{Cq Lysozyme Midgut}$ | .778 | .074 | -.092 |
| $\Delta\Delta\text{Cq GST1 Fatbody}$ | -.473 | .611 | -.008 |
| $\Delta\Delta\text{Cq GST1 Midgut}$ | -.032 | .724 | -.007 |
| $\Delta\Delta\text{Cq Transferrin Fatbody}$ | .627 | -.127 | .533 |
| $\Delta\Delta\text{Cq Transferrin Midgut}$ | .210 | .000 | .741 |
| $\Delta\Delta\text{Cq iNOS Fatbody}$ | .386 | -.523 | .115 |
| $\Delta\Delta\text{Cq iNOS Midgut}$ | .062 | .454 | .046 |

Extraction Method: Principal Component Analysis. Rotation Method: Oblimin with Kaiser Normalization.

